# Common cuckoo female host selection is not determined by host quality but can affect cuckoo nestling growth when parasitising Common redstarts

**DOI:** 10.1101/2024.03.11.583276

**Authors:** Teresa Abaurrea, Angela Moreras, Jere Tolvanen, Robert L. Thomson, Rose Thorogood

## Abstract

Common cuckoo (*Cuculus canorus*) females lay their eggs in the nests of other avian hosts, relying on parental care provided by parasitised hosts. Therefore, it would benefit cuckoo females to target high-quality individual hosts, able to provide optimal parental care. Attempts at testing cuckoo female host selection have so far shown mixed results. However, this might be because studies have rarely considered the host nests that are available in space and time during each cuckoo egg-laying event, as well as the implications of host choice on the cuckoo nestling growth. Here we combined long-term monitoring data with an experiment to test whether cuckoo females parasitising Common redstarts (*Phoenicurus phoenicurus*) target individual hosts to optimise their nestlingś growth. Making use of data collected between 2013 and 2022 from 350+ nest boxes in Oulu (Finland), we first explored whether there is a range of available host nests for the cuckoo female to choose from. Second, we tested if hosts are targeted according to individual quality (using clutch size as a proxy). And third, we investigated the outcomes of cuckoo female host-selection on nestling growth (mass, tarsus length, and wing length) between 2014 and 2019. We conducted a cross-fostering experiment where we either left cuckoo eggs to hatch and be raised in the nest their mother originally chose for them or moved cuckoo eggs to non-parasitised nests. Additionally, we conducted an exploratory analysis to test the quality of the parents caring for the cross-fostered cuckoo nestlings. After accounting for how many host nests were available to the cuckoo female to choose from, we found that nests with bigger clutches were not more likely to be parasitised, and cross-fostering did not affect mass and tarsus length growth. However, both in the wing length growth and in our exploratory analysis of host parental care on cuckoo nestling growth we found that cuckoo nestlings that grew in the nest selected by their mothers, reached higher asymptotic growth at a slower rate. This suggests that while cuckoos may not choose redstart hosts based on their individual quality when parasitising common redstarts, cuckoo female host selection might improve cuckoo nestling growth and thus, have an adaptive significance.

## INTRODUCTION

Most bird species invest in parental care beyond egg production, however, avian brood parasites avoid the cost of providing parental care to their young by laying their eggs in the nests of other avian species (Davies, 2000). Once the eggs hatch, brood parasite nestlings rely on their host parentś care (Davies, 2000), which can vary with individual quality. High-quality parents are able to select territories with higher abundance (Decker et al., 2012) and/or quality of food resources (Wilkin et al., 2009). This implies that high-quality parents can provision more food of better quality to nestlings (Pagani-Núñez & Senar, 2014), resulting in measurable effects that improve nestling growth and survival (Schwagmeyer & Mock, 2008; Silva et al., 2007). Therefore, it should be adaptive for parasitic females to lay their eggs in nests of high-quality host parents to increase the chances of their offspring fledging in good condition (Louder et al., 2015). However, despite studies exploring this across different host species, there is surprisingly little empirical support for brood parasites assessing individual quality of hosts and adjusting their laying decisions accordingly.

Evidence showing that brood parasite females select nests with high-quality hosts is not conclusive. Research suggests that brood parasites select high-quality hosts based on different individual quality proxies such as female host body condition (Polačiková et al., 2009), male host alarm calling behaviour (Marton et al., 2019), host egg volume (Alvarez, 2000), nest building activity (Banks & Martin, 2001), or nest size (Soler et al., 1995). It has also been shown that host quality is important only when brood parasitism rate is low (Molina-Morales et al., 2016). Alternatively, rather than selecting host nests based on the potential for high-quality care, parasitic females might make oviposition decisions based on the appearance of the host’s own eggs (i.e. egg-matching strategy to decrease egg-rejection by hosts, Zhang et al., 2023). Or oviposition decisions may be driven more by variation in nest detectability (e.g., nest visibility in great reed warblers, Moskát and Honza, 2000) rather than host quality (although these may be correlated, Stokke *et al*., 2008).

To choose high quality individuals, brood parasite females require multiple host nests to be available (Jelínek et al., 2014). However, studies exploring host choice did not account for temporal and spatial availability of host nests (Lyon, 1993). In almost all brood parasite – host systems, the parasitism of host nests can only occur within the home range of the parasite female, and within a short temporal window of suitability. For example, Common cuckoo females (*Cuculus canorus*, cuckoo hereafter) only lay eggs within their territory and usually only while the host is still egg-laying (Davies, 2000; Koleček et al., 2021). Therefore, to be able to detect patterns of oviposition depending on host quality, it is critical that the scope for choice needs to be considered (Lyon, 1993).

In addition to overlooking host availability when studying host selection, the effect of any potential femalés choice on her offspring growth and survival has been neglected, which impairs our understanding of whether these choices are adaptive (Marshall & Uller, 2007). There is a growing body of knowledge on how brood parasite nestling growth varies depending on host care. It has been shown that growth of parasite chicks can vary between different species of hosts (Grim, 2006; Grim et al., 2014; Winnicki et al., 2021), even if these have the same breeding ecology (Grim & Samaš, 2016). Furthermore, brood parasitic nestling growth can also vary depending on whether the parasite grows up alone, along host nestlings (Geltsch et al., 2012), or even depending on the social mating status of its host (Trnka et al., 2012). However, to our knowledge, there are no studies on the effect of host individual quality on the parasitic nestling growth. An alternative approach to better test for individual host choice would be to experimentally compare the outcomes of disrupting that choice.

Here we used a nest-box breeding study population of Common redstarts (*Phoenicurus phoenicurus*, redstarts hereafter) that has been regularly parasitised by the Common cuckoo (*Cuculus canorus*). Cuckoo eggs at our study site are only laid by redstart-gens (i.e. immaculate blue egg) cuckoo females which are highly host specific to redstart nests as their blue eggs are excellent mimics of this host’s eggs (Fossøy et al., 2016). These two characteristics (nest-box breeding population and high brood parasite specificity) provides a unique opportunity to conduct brood parasitism studies accounting for the majority of potential cuckoo host nests in the area.

Our main aim was to test whether cuckoo females target host nests to optimise their nestlingś growth accounting for nest availability, using a readily accessible host quality proxy and manipulating the femalés host choice outcome. First, we explored whether there is a range of available host nests to choose from. Second, we used a georeferenced 9-year-long breeding monitoring dataset to describe spatio-temporal patterns of host nest availability and parasitism occurrence according to clutch size. We used clutch size as a proxy because it is a reliable indicator of parental investment (Garamszegi et al., 2004; Houston et al., 1983) and individual quality in other avian species (Slagsvold & Lifjeld, 1990). We predicted that high-quality redstart individuals laying larger clutches would be more likely to be parasitised than other redstart host nests available within the cuckoo’s spatio- temporal laying area. Third, we conducted a cross-fostering experiment to test whether host nest choice affects the growth of cuckoo nestlings (common proxy of fitness used in studies of birds, e.g., Bryant, 1978). We predicted that cuckoo nestlings raised in the host nest chosen by their mother would grow larger and/or faster than cuckoo nestlings moved to non-parasitised redstart nests.

## METHODS

### Study site and nest monitoring

Fieldwork was conducted in the forests around the city of Oulu, Finland (64°60’N, 25°42’E). The forest patches were predominantly Scots pines (*Pinus sylvestris*) with low vegetation ground cover of lingonberry (*Vaccinium vitis-idaea*), blueberry (*Vaccinium myrtillus*), heather (*Calluna vulgaris*), mosses (*Pleurozium schreberi* and *Dicranum spp*) and lichens (*Cladonia spp*) (Rutila et al., 2002).

Since 2002, commercially-available wooden nest boxes (Linnunpönttö Oy) have been placed on pines at approximately 1.5 m above the ground and separated by 100 – 250 m in an area of approximately 60 km^2^ (Thomson et al., 2016). The boxes measure 17.5 x 17.5 x 28 cm (width x depth x height) with a 7 cm diameter entrance hole and are emptied each spring (except in 2020) before the first redstarts and cuckoos arrive at the study area in early May, after their migration from sub- Saharan Africa. During our study, taking place between 2013 and 2022 (except 2020), the number of nest boxes increased from 271 to 374, and all were georeferenced using hand-held GPS (± 5 m accuracy).

Nest boxes were monitored for breeding activity at least once a week throughout the breeding season and we identified the occupying species through visual inspection of nest materials, egg colour and pattern, and nearby adults. Once a nest box was occupied by redstarts (the majority of occupants, see Results), it was monitored approximately every 3 – 5 days to record laying date, clutch size, incubation status, and presence of any cuckoo eggs. If the laying of the first egg was missed, then the date of clutch initiation was back-calculated assuming that redstart females lay one egg per day during the laying period (Thomson et al., 2016). The expected hatching date for cuckoo eggs was estimated by calculating 12 days from the onset of incubation. Nests were monitored until predation or abandonment occurred, or until nestlings were close to fledging to estimate fledging success (redstart nestlings: 15 days, cuckoo nestlings: 21 days). Whenever a nest box was visited, we approached carefully to check for any possible mislaid cuckoo eggs outside the box (i.e. eggs on the ground, possibly from a failed cuckoo laying attempt since redstarts rarely eject naturally laid cuckoo eggs to avoid underestimating parasitism, see (Thomson et al., 2016) for more details).

### Cross-fostering experimental design and protocol

To test whether a cuckoo’s choice of host nest may be adaptive, we conducted a cross-fostering experiment with cuckoo eggs laid between 2014 and 2019. We randomly assigned cuckoo eggs to one of two treatments: (i) ‘chosen’ – the cuckoo egg was left to hatch in the redstart nest as intended by the cuckoo female; (ii) ‘non-chosen’ – the cuckoo egg was relocated to a non-parasitised redstart nest nearby. In the non-chosen treatment, cuckoo eggs were moved as soon as detected during routine monitoring of nests at the egg-laying stage. To standardise the manipulation of eggs in both treatments, eggs that were left to hatch in their original nests (i.e. chosen treatment) were removed momentarily and returned to their nest. During this experiment, we recorded location and phenology information of both the nest where each cuckoo egg was laid and its destination nest after cross- fostering. After hatching, cuckoo nestlings were measured at 3, 6, 9, 12, 15 and 18 days of age to record their weight (to the nearest 0.01 g), tarsus length (to the nearest 0.01 mm), and wing length (to the nearest millimetre) – for logistical reasons, some nestlings were not measured at all ages.

### Data analysis methods

All data analyses detailed below was conducted in R version 4.3.2 (2023-10-31 ucrt, R Core Team, 2023).

### Spatio-temporal variation in parasitism

To estimate spatio-temporal variation of parasitism in our study area, we built a generalised additive mixed model (GAMM) implemented using package *mgcv* (version 1.9.0, Wood, 2017). These GAMMs provide enough flexibility to include non-linear predictive variables and account for spatial and temporal autocorrelation. Parasitism occurrence was modelled using a binomial distribution and logit link function, with laying date included as a smoother. We also included year as random effect, and geographical coordinates as a 2-dimensional full tensor product smoother, suitable for covariates in different scales or units, such as latitude and longitude (Zuur, 2014).

### Host nest availability

Although redstart-cuckoo gentes only parasitise redstart nests, redstart nests are not always available for parasitism if the clutch has already been completed (mean laying period ± SD = 6.67 ± 0.82 days), or if they are built beyond a cuckoo’s breeding territory. As we did not phenotype or genotype eggs to cuckoo females during this study, we instead made assumptions about which nests were available for parasitism based on the most comprehensive published estimates of the size of cuckoo females’ breeding areas available (Koleček et al., 2021). We first calculated active host nest density in our study area (median = 0.09 nests/ha/year); as this was much lower than the breeding density of great reed warblers parasitised by common cuckoos reported by Koleček *et al*. (median = 0.29 nests/ha, Koleček *et al*., 2021), we extrapolated a biologically-informed estimate of cuckoos’ breeding area in our study area accordingly (Koleček *et al*., 2021: 65 ha; our study area: 195 ha).

Redstart nest boxes are distributed radially throughout the forest, so we next tabulated the number of all redstart nests available for parasitism (i.e. still laying eggs) in time and space using an 800- meter radius circle centred on each nest. For analyses of parasitism occurrence and clutch size, we only included nests with two or more alternative available nests to ensure that female cuckoos had a choice. To guarantee the robustness of our conclusions, we also calculated nest availability using a smaller (400 m radius) and a larger (1200 m radius) putative cuckoo breeding area (see supplementary materials).

### Spatio-temporal variation in redstart clutch size

Before analysing the relationship between host clutch size and parasitism occurrence, we checked for spatio-temporal patterns in redstart clutch size, since clutch sizes can decrease with laying date (Gladbach et al., 2010) and can be influenced by local conditions (Hussell, 1972). We built a generalised additive mixed model (GAMM) with Gaussian distribution and identity link function and modelled clutch size as a function of laying date as a smoother, year as random effect, and the geographical coordinates as a 2-dimensional full tensor product smoother. As expected, clutch size and laying date were negatively correlated [GAMM: s(Laying date, 2.61), Ref.df = 9.0, F = 9.86, p- value < 0.001, R2(adj) = 0.09, Figure 1A]. We also found that nests in the northern parts of our area had significantly larger clutch sizes [GAMM: te(Longitude, Latitude, 2.11), Ref.df =23.0, F = 0.27, p- value = 0.029, R2(adj) = 0.09, Figure 1B]. To control for this spatio-temporal variation in clutch size, we used Pearson residuals from the GAMM (i.e. difference between observed clutch size and the expected clutch size for a given date and location) rounded to the nearest egg number (i.e. spatio- temporally adjusted clutch size, STA clutch size hereafter, Figure 1C) in all further analyses.

**Figure 1:**
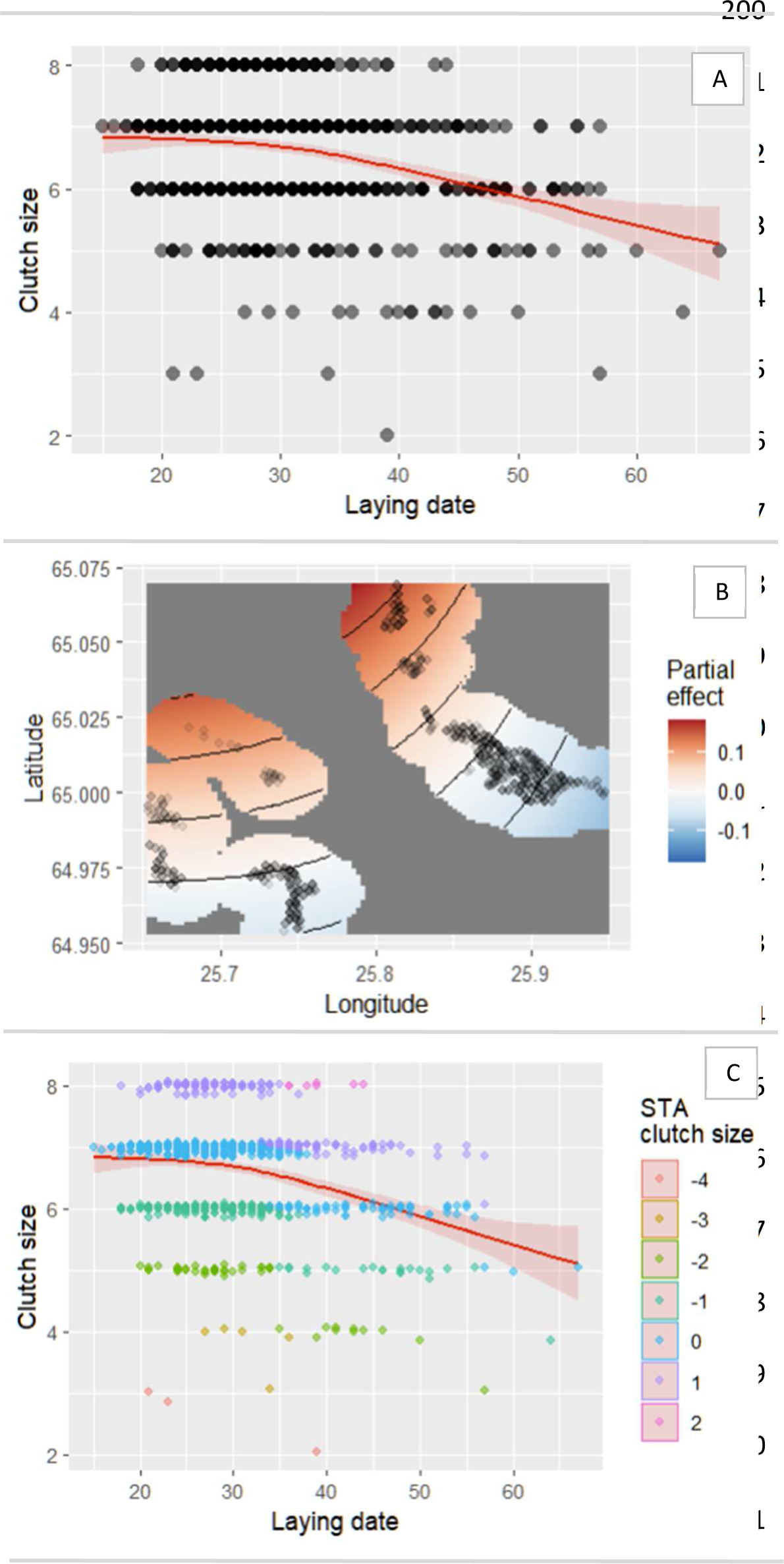
Results of the GAMM analysing clutch size and used to create the spatio-temporally adjusted clutch size variable (STA clutch size). **A)** Raw data plot of clutch size *versus* laying date, with the GAMM (in red) that indicates that clutch size decreases throughout the season. Shaded area represents 95% confidence intervals. **B)** Spatial distribution of clutch size data in our study area showing a gradient decreasing from north-west (red) to south-east (blue). **C)** STA clutch size continuous variable based on the GAMM (in red). Here, STA clutch size is transformed into categorical variable with different categories in different colours to facilitate visualisation (from -4 to 2, where -4 indicates nests that are 4 eggs smaller than expected for a specific date, and 2 indicates nests that are 2 eggs larger than expected for a specific date).

### Parasitism occurrence according to host clutch size

To test whether high quality hosts, with bigger clutches, are more likely to be parasitised, we built a generalised linear mixed model (GLMM) by Laplace approximation with binomial distribution and logit link to model parasitism as a function of STA clutch size and number of available nests within the assumed breeding area of a local cuckoo as predictive variables (using R package *lme4*, version 1.1- 35.1, Bates *et al*., 2014). We included year as a random effect to account for environmental variation and changes in samples size. Using Akaikés Information Criterion value we compared this main model with the following models: a model that included only random effect year, a model including STA clutch size and year as random effect, and a model accounting for the interactions between STA clutch size and the number of available nests, and year as random effect.

For these analyses, the dataset included only nest boxes occupied by redstarts with complete information available for redstart laying date, clutch size, and latitude and longitude, and where we were confident that our monitoring did not miss possible parasitism events (n = 961). This filtering removed nests that were abandoned before or during laying (redstarts usually start building several nests before completing one, pers. obs.), or depredated before clutch completion. The dataset was then truncated according to when the first and last cuckoo egg of the season was laid (in 2013, n = 1; in 2015, n = 1; 2019, n = 4; in 2021, n = 1, and in 2022, n = 2) to limit host nests to those available for parasitism. Finally, any nests with clutches larger than 9 eggs (n = 7) were excluded from the analyses because these larger clutches are usually laid by multiple females (pers. obs. – most likely because a female takes over an occupied and incubating nest and starts egg laying without building a new nest cup). We then excluded parasitised nests with no cuckoo egg laying date (n = 5), where we were uncertain of the cuckoo egg laying date by more than 5 days (n = 31), and those where the cuckoo egg was laid outside the redstart egg laying period (n = 20). After accounting for spatio-temporal nest availability (remove nests with only one nest available, n = 86), the final sample size used in this analysis was 803 nests.

### Effects of host nest choice on cuckoo nestling growth

We estimated the asymptote and growth rate of weight, tarsus length and wing length by fitting a standard logistic growth model (Ricklefs, 1968) with an SSlogis function in R (*stats* package, R Core Team, 2023). To our knowledge there is no study analysing the best growth model fit in Cuculiformes, but it has been shown that the logistic growth model is flexible enough to fit well multiple avian orders (Passeriformes (Starck & Ricklefs, 1998), Galliformes (Aggrey, 2002) and Columbiformes (Gao et al., 2016)). We calculated these growth parameters for 40 cuckoo nestlings but removed those that died before 18 days of age (n = 4 from three different years, which growth curves decreased sharply suggesting lack of food or an underlying condition). Additionally, there were three nestlings where wing length was measured only three times, not enough timepoints to fit the growth model.

Thus, wing models included data on 33 nestlings only.

Here, we built GAMMs with Gaussian distribution and identity link function to include non-linear predictive variables in the models, such as hatching date. We used the asymptote and growth rate of weight, tarsus length, and wing length as response variables. These models included the type of nest as a parametric predictor, with a well-balanced sample size (18 nestlings in chosen nests, and 18 in non-chosen nests). We also incorporated hatching date as a smoother, and year as random effect to control for different sample sizes between years and different environmental conditions. These models also controlled for the number of available nests (see previous section for definition of number of available nests) as weights, defined here as *Available nests/mean(Available nests)*.

In addition to these models, we conducted an exploratory analysis that accounted for the quality of the parents caring for the cross-fostered cuckoo nestlings. Here, we only included nestlings that were moved to non-chosen nests (n = 18) to analyse the effect of being fostered by parents of lower, similar, or higher quality than the ones originally chosen by the cuckoo female on the cuckoo nestling growth. We built linear models with clutch size difference (i.e., the numeric difference between the redstart clutch size in the cuckoo nestlinǵs original nest and the clutch size in the nest where the cuckoo hatched) as polynomial explanatory variable.

### Ethical note

This study was conducted following the current laws and recommendations for animal welfare of Finland. Permits to place nest boxes in our study area were granted to Jere Tolvanen by the City of Oulu (564-2017-8 and 564-2020-4). Nest monitoring until 2019, and the cross-fostering experiment was conducted under research permits POPELY/136/07.01/2014 and VARELY/921/2017, granted by Elinkeino-, liikenne- ja ympäristökeskus (Centre for Economic Development, Transport, and the Environment of Finland) to Jere Tolvanen. In 2021 and 2022, nest monitoring research permit (ESAVI/12343/2020) was granted also by Elinkeino-, liikenne- ja ympäristökeskus to Teresa Abaurrea.

## RESULTS

### Cuckoo female selection of host nests

During the study, redstarts occupied over half of the nest boxes provided each year (average N per year ± SD = 163 ± 33, range = 120 – 219, which represents 67 ± 5 % per year). Five other species were also observed using the nest boxes (great tits (average N per year ± SD = 47 ± 16, range = 10 – 62, 19 ± 6 % per year), pied flycatchers (average N per year ± SD = 20 ± 9, range = 12 – 41, 9 ± 4 % per year), crested tits (*Lophophanes cristatus*, 4 nests occurring in 3 years), blue tits (*Cyanistes caeruleus*, 1 nest), and willow tits (*Poecile montanus*, 1 nest), but none of these nests were parasitised.

In our area, peak egg-laying for redstarts occurred on the 27^th^ May, while cuckoos showed two peaks of parasitism, 27^th^ May and 16^th^ June. These peaks were estimated using a function that calculated all peaks of a histogram built with Sturges method. The function was coded using package *base* (R Core Team, 2023). Furthermore, we found that the risk of parasitism was not constant during the breeding seasons [GAMM: s(Laying date, 1.95), Ref.df = 9.0, χ² = 54.56, p-value < 0.001, R2(adj) = 0.09] or across the study site [GAMM: te(Lon, Lat, 2.19), Ref.df = 24.0, χ² = 6.64, p-value = 0.017, R2(adj) = 0.09, Figure 2], with some areas parasitised less often than others (Figure 2).

**Figure 2:**
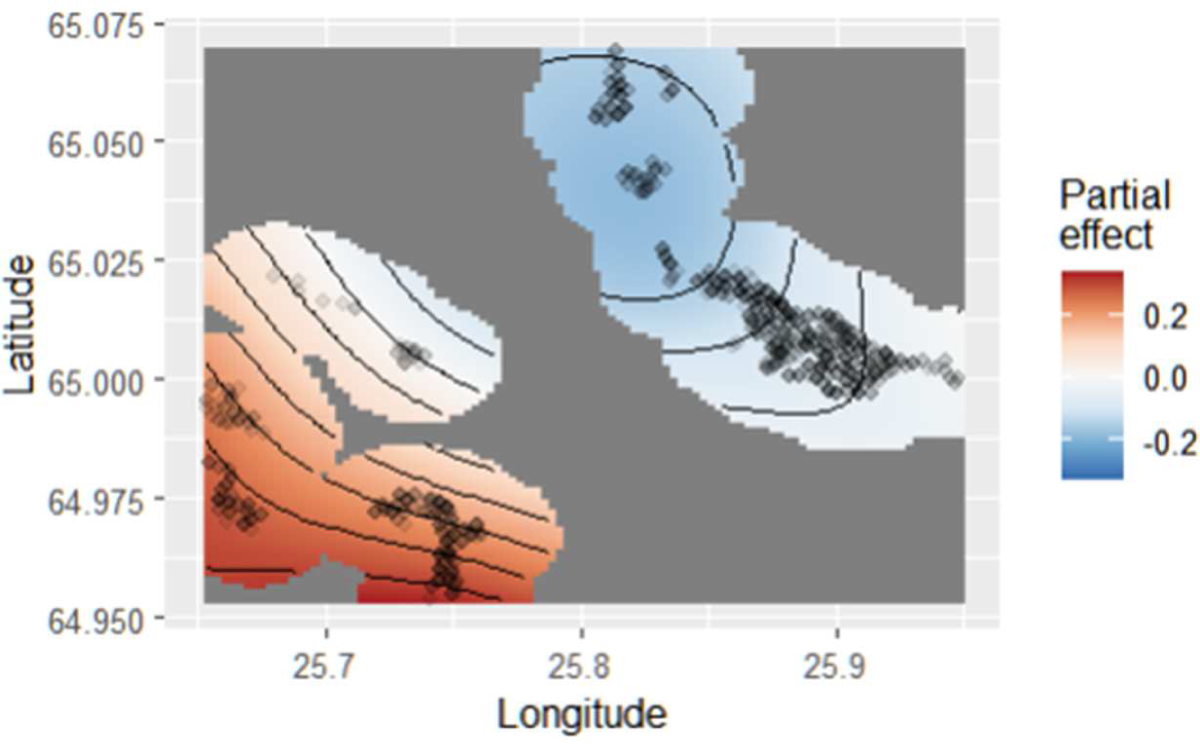
Map of parasitism occurrence estimated from the generalised additive mixed model and showing the partial effects of geographical coordinates within our study area.

In total, we followed 223 nests that were parasitised by a cuckoo. On average, 24% of redstart nests were parasitised at least once by cuckoos (average N parasitised redstart nests per year ± SD = 25 ± 12, range = 9 – 43 nests). Due to multiple parasitism events per nest, the average N of cuckoo eggs per year was 28 ± 14 (range = 9 – 50). Within our estimated cuckoo female breeding area, there was only one redstart nest available for the cuckoo female on 86 occasions (9.7 %), and thus no choice. When a choice was possible, there were between 2 and 24 nests available for the cuckoo female to choose from, with four nests available as the most frequent situation (Figure 3). Most clutches (81.3% of 803) had 6 or 7 eggs, and these two clutch sizes experienced most parasitism events (76.7% of 189) (Table 1).

**Figure 3:**
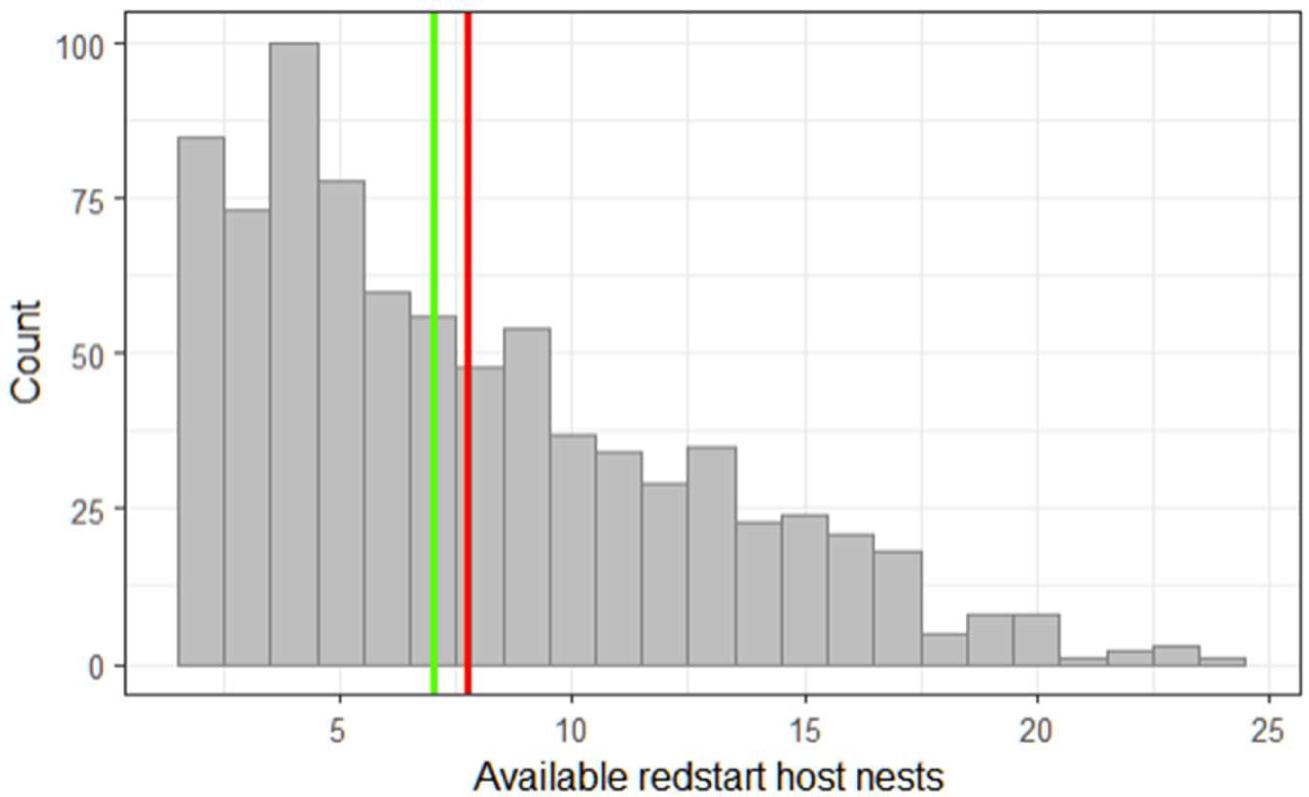
Histogram of number of available nests during each parasitism event, where the green vertical line indicates the median (7 nests), and the red line indicates the mean (7.76 nests).

**Table 1.** Availability of redstart nests and parasitism occurrence according to clutch size (percentage of total nests in parentheses).

| *Clutch size* | *Number of redstart nests* | *Parasitised nests* | *Proportion of nests parasitised* |
| --- | --- | --- | --- |
| 2 | 1 (0.1%) | 0 (0.0%) | 0/1 (0.0%) |
| 3 | 1 (0.1%) | 0 (0.0%) | 0/1 (0.0%) |
| 4 | 9 (1.1%) | 4 (2.1%) | 4/9 (44.4%) |
| 5 | 40 (5.0%) | 15 (7.9%) | 15/40 (37.5%) |
| 6 | 251 (31.3%) | 64 (33.9%) | 64/251 (25.5%) |
| 7 | 402 (50.1%) | 81 (42.9%) | 81/402 (20.1%) |
| 8 | 99 (12.3%) | 25 (13.2%) | 25/99 (25.3%) |
| *TOTAL NESTS* | 803 | 189 |  |

Our model selection process indicated that the best fitting model included STA clutch size and number of available nests with no interaction, as well as year as random effect (i.e., our initial model, Table 2). Contrary to our expectations, however, the probability of being parasitised did not increase with clutch size, and instead only correlated significantly with the number of nests available for a cuckoo to choose from – when there were more nests available, parasitism probability of individual nests declined (Figure 4 and Table 3). These results were consistent using a 400 m (Intercept = -0.43; STA clutch size estimate = -0.09, p-value = 0.40; Available nests estimate = -0.27, p-value < 0.001) and 1200 m radius (Intercept = -0.48; STA clutch size estimate = -0.09, p-value = 0.36; Available nests estimate = -0.07, p-value < 0.001) for cuckoo female breeding areas.

**Figure 4:**
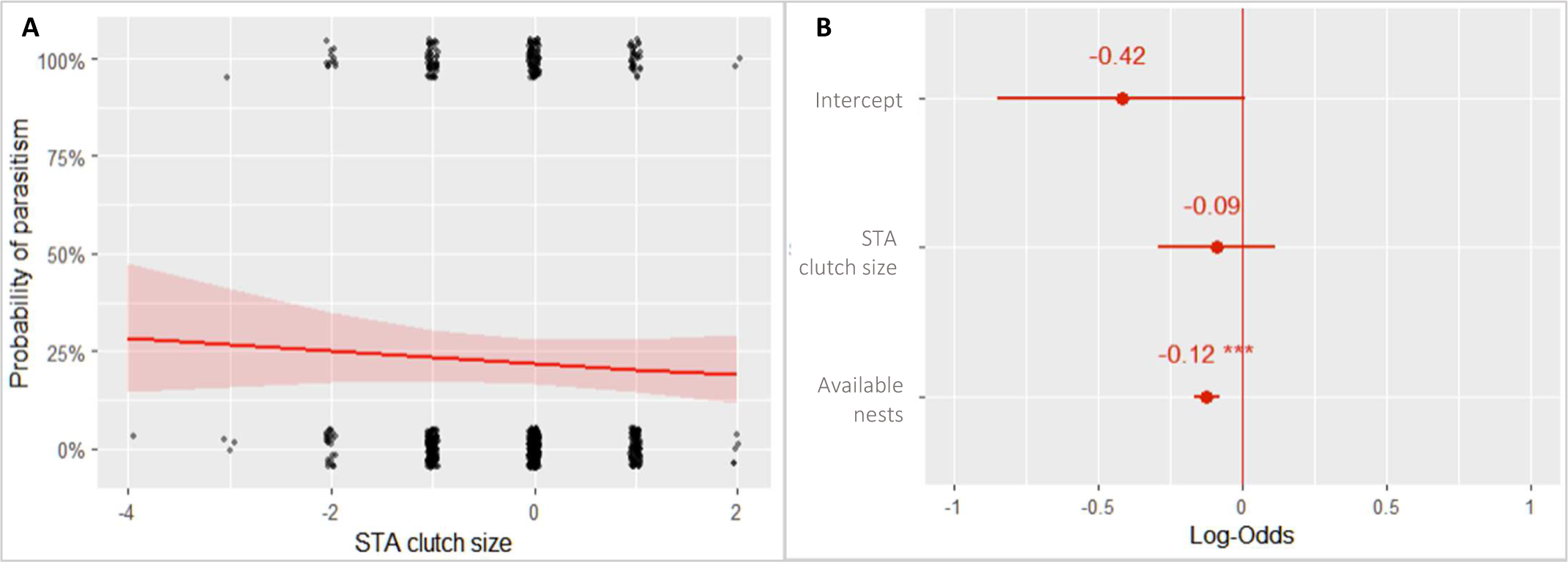
Relationship between probability of parasitism and STA clutch size. **A)** The probability of parasitism shows a non-significant decrease with spatio-temporally adjusted (STA) clutch size (where negative clutch sizes indicate smaller than expected clutches and positive clutch sizes indicate larger than expected clutches). The plot shows the relationship between raw data points, and the shaded band indicates 95% confidence interval. To increase visibility, data points are jittered. **B)** Summary results of GLMM relating the probability of parasitism to predictive variables included in the model (STA clutch size and number of available nests). Values represented are estimated model coefficients (mean ± 95% confidence interval) and respective significance (*** , P<0.001; ** , P<0.01; * , P<0.05; . , P<0.1).

**Table 2.** Model selection of parasitism GLMM.

| *Model* | *K* | *AIC* | *Delta AIC* | *AIC weight* |
| --- | --- | --- | --- | --- |
| ~ STA clutch size + Available nests + (1 \| Year) | 4 | 837.34 | 0.00 | 0.73 |
| ~ STA clutch size * Available nests + (1 \| Year) | 5 | 839.31 | 1.97 | 0.27 |
| ~ 1 + (1 \| Year) | 2 | 868.70 | 31.36 | 0.00 |
| ~ STA clutch size + (1 \| Year) | 3 | 870.01 | 32.67 | 0.00 |

**Table 3.**
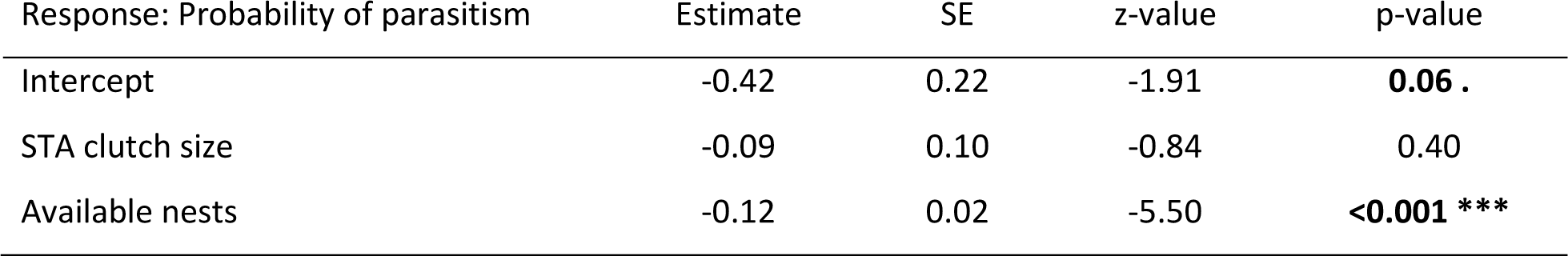
GLMM explaining the parasitism probability of redstart nests according to spatio-temporally adjusted (STA) clutch size. Number of available nests is included as a covariate to control for variation in potential choices. SE: standard error. Significant p-values in bold (*** , P<0.001; ** , P<0.01; * , P<0.05; . , P<0.1).

### Consequences of cuckoo female choice on cuckoo nestling growth

Models including type of cross-fostered nest as a predictor variable showed no significant differences in weight or tarsus length growth parameters (asymptote and growth rate; Table 4 and Figure 5-1-ii and 5-2-ii). However, cuckoo nestlings that grew in the nest chosen for them by their mother had marginally significantly longer wings that grew at a marginally non-significantly slower rate than nestlings cross-fostered to a different nest (Table 4 and Figure 5-3-ii).

**Figure 5:**
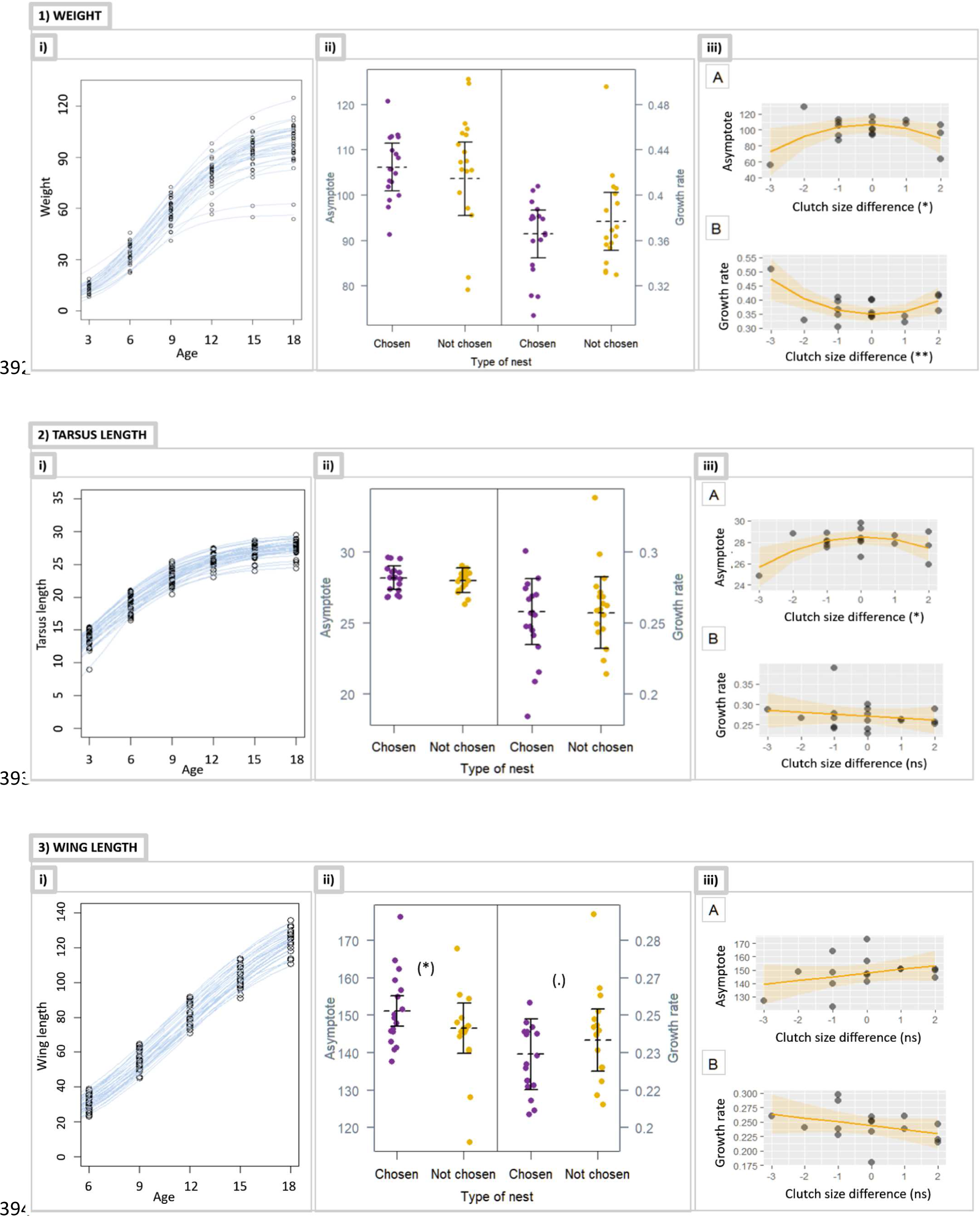
Model plots for weight, tarsus length and wing length results. Within each morphometric (1, weight, 2, tarsus length, and 3, wing length), plots labelled with “i” are growth curves of all nestlings (n = 36 for weight and tarsus length, n = 33 for wing length). Plots labelled with “ii” show models with response variables (asymptote and growth rate for all nestlings - n = 36 for weight and tarsus length, n = 33 for wing length) as a function of two covariates (type of nest, hatching date) and one random effect (year). Plots labelled with “iii” show models for the asymptote (A) and growth rate (B) as a function of clutch size difference (n = 18 for weight and tarsus length, n = 16 for wing length).

**Table 4.**
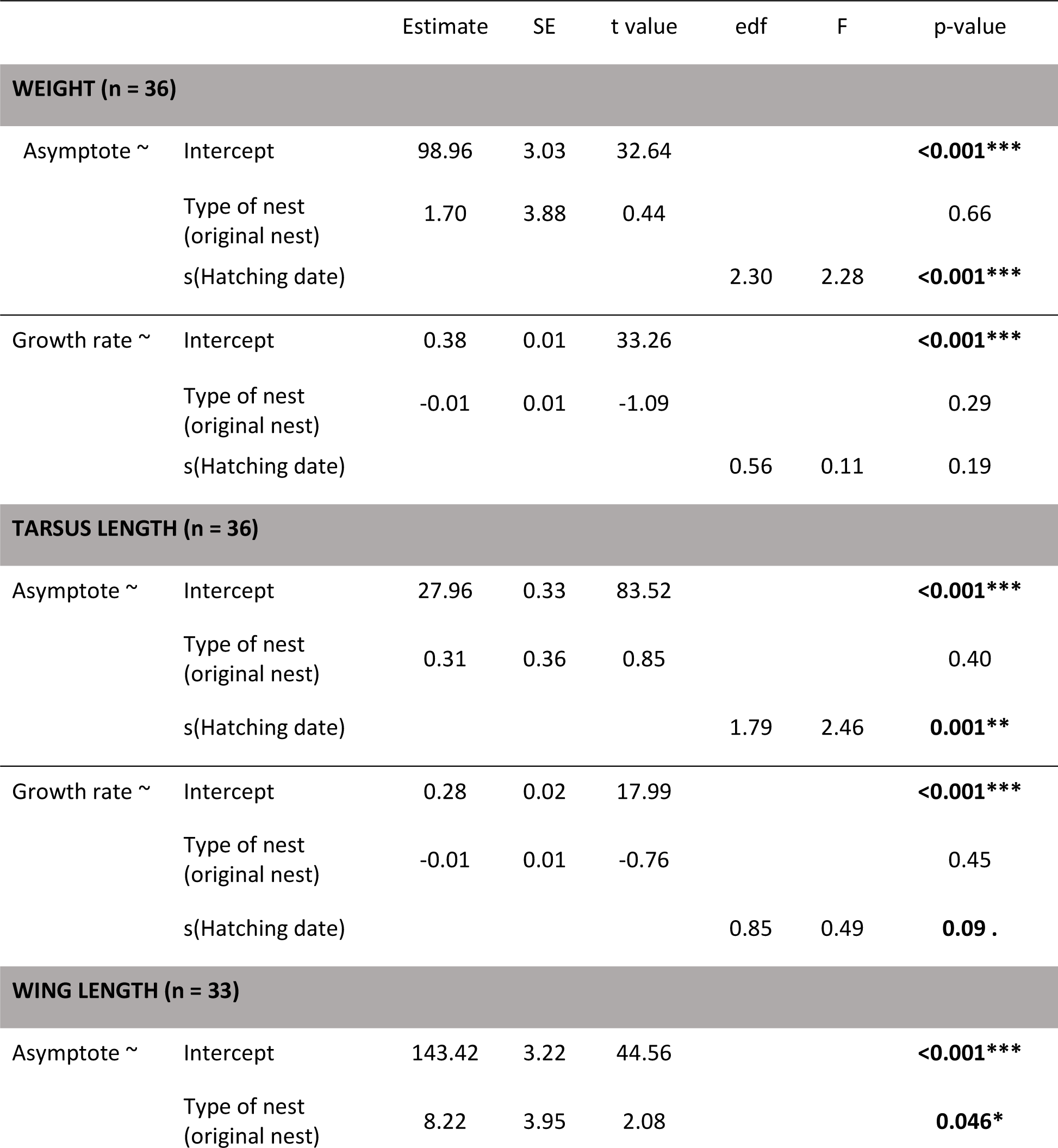
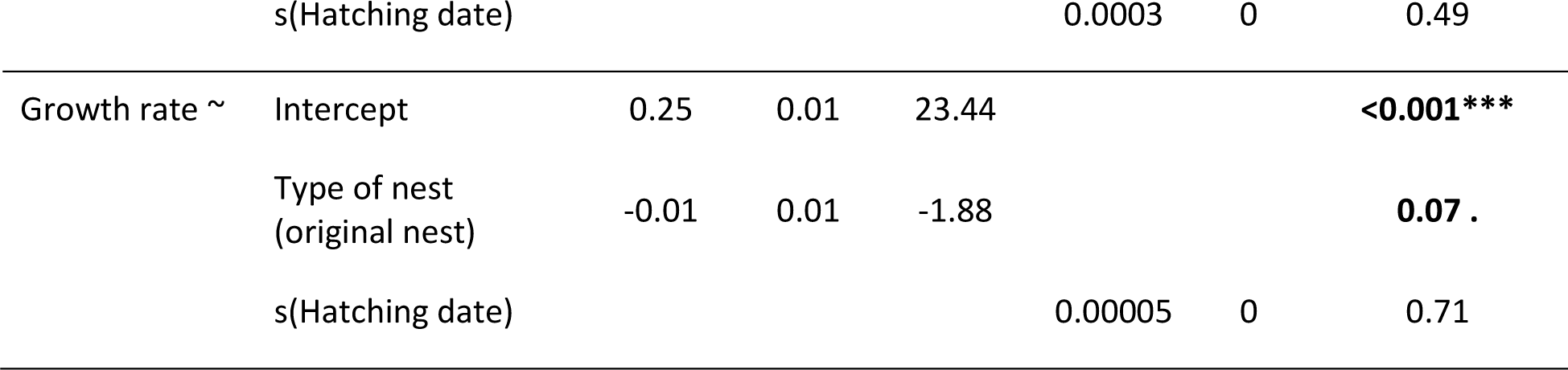
Summary statistics of the GAMMs (Gaussian distribution with identity link) explaining the different cuckoo nestling morphometrics, and including type of nest as predictive variable. Significant p-values in bold (*** , P<0.001; ** , P<0.01; * , P<0.05; . , P<0.1). SE, standard error; t value, T statistic; edf, estimated degrees of freedom; F, F statistic.

|  |  | Estimate | SE | t value | edf | F | p-value |
| --- | --- | --- | --- | --- | --- | --- | --- |
| **WEIGHT (n = 36)** |  |  |  |  |  |  |  |
| Asymptote ~ | Intercept | 98.96 | 3.03 | 32.64 |  |  | **<0.001\*\*\*** |
|  | Type of nest<br>(original nest) | 1.70 | 3.88 | 0.44 |  |  | 0.66 |
|  | s(Hatching date) |  |  |  | 2.30 | 2.28 | **<0.001\*\*\*** |
| Growth rate ~ | Intercept | 0.38 | 0.01 | 33.26 |  |  | **<0.001\*\*\*** |
|  | Type of nest<br>(original nest) | -0.01 | 0.01 | -1.09 |  |  | 0.29 |
|  | s(Hatching date) |  |  |  | 0.56 | 0.11 | 0.19 |
| **TARSUS LENGTH (n = 36)** |  |  |  |  |  |  |  |
| Asymptote ~ | Intercept | 27.96 | 0.33 | 83.52 |  |  | **<0.001\*\*\*** |
|  | Type of nest<br>(original nest) | 0.31 | 0.36 | 0.85 |  |  | 0.40 |
|  | s(Hatching date) |  |  |  | 1.79 | 2.46 | **0.001\*\*** |
| Growth rate ~ | Intercept | 0.28 | 0.02 | 17.99 |  |  | **<0.001\*\*\*** |
|  | Type of nest<br>(original nest) | -0.01 | 0.01 | -0.76 |  |  | 0.45 |
|  | s(Hatching date) |  |  |  | 0.85 | 0.49 | **0.09 .** |
| **WING LENGTH (n = 33)** |  |  |  |  |  |  |  |
| Asymptote ~ | Intercept | 143.42 | 3.22 | 44.56 |  |  | **<0.001\*\*\*** |
|  | Type of nest<br>(original nest) | 8.22 | 3.95 | 2.08 |  |  | **0.046\*** |
|  | s(Hatching date) |  |  |  | 0.0003 | 0 | 0.49 |
| Growth rate ~ | Intercept | 0.25 | 0.01 | 23.44 |  |  | **<0.001\*\*\*** |
|  | Type of nest<br>(original nest) | -0.01 | 0.01 | -1.88 |  |  | **0.07 .** |
|  | s(Hatching date) |  |  |  | 0.00005 | 0 | 0.71 |

To explore these results further, we investigated whether the difference in clutch size between the nest of origin and the cross-fostered destination nest could explain differences in nestling growth. We found that cuckoo nestlings moved to nests with a host clutch size similar to their original nest tended to grow heavier (asymptotic weight, Figure 5-1-iii-A) albeit more slowly (weight growth rate, Figure 5-1-iii-B) than those moved to nests with a smaller clutch size (Table 5). These nestlings also grew larger in terms of tarsus length (asymptote, Figure 5-2-iii-A) but not for wing length (asymptote, Figure 5-3-iii-A). There were no differences in the growth rate of these morphometrics (Figure 5-2-iii- B, 5-3-iii-B, Table 5). However, these results were influenced strongly by one cuckoo nestling moved to a host nest with a much smaller clutch size (Asymptotic weight: Intercept = 102.27, CS difference estimate = -24.1, p-value = 0.11; CS difference^2 estimate = 2.17, p-value = 0.88; Weight growth rate: Intercept = 0.36, CS difference estimate = 0.04, p-value = 0.24; CS difference^2 estimate = 0.02, p-value = 0.66; Asymptotic tarsus length: Intercept = 28.15, CS difference estimate = -0.91, p-value = 0.36; CS difference^2 estimate = -0.47, p-value = 0.63) and so should be interpreted with caution.

**Table 5.** Summary statistics of the LMs explaining the different cuckoo nestling growth parameters, and including clutch size difference as predictive variable. Significant p-values in bold (*** , P<0.001; ** , P<0.01; * , P<0.05; . , P<0.1). SE, standard error; t value, T statistic.

|  |  | Estimate | SE | t value | p-value |
| --- | --- | --- | --- | --- | --- |
| **WEIGHT (n = 18)** |  |  |  |  |  |
| Asymptote ~ | Intercept | 99.71 | 3.75 | 26.62 | **<0.001\*\*\*** |
|  | Clutch size difference | 3.14 | 15.89 | 0.20 | 0.85 |
|  | Clutch size difference ^2 | -38.95 | 15.89 | -2.45 | **0.027\*** |
| Growth rate ~ | Intercept | 0.37 | 0.01 | 39.53 | **<0.001\*\*\*** |
|  | Clutch size difference | -0.04 | 0.04 | -0.93 | 0.37 |
|  | Clutch size difference ^2 | 0.12 | 0.04 | 3.13 | **0.007\*\*** |
| **TARSUS LENGTH (n = 18)** |  |  |  |  |  |
| Asymptote ~ | Intercept | 27.97 | 0.24 | 117.24 | **<0.001\*\*\*** |
|  | Clutch size difference | 0.90 | 1.01 | 0.89 | 0.39 |
|  | Clutch size difference ^2 | -2.81 | 1.01 | -2.78 | **0.01\*** |
| Growth rate ~ | Intercept | 0.27 | 0.01 | 31.67 | **<0.001\*\*\*** |
|  | Clutch size difference | -0.005 | 0.01 | -0.76 | 0.46 |
| **WING LENGTH (n = 16)** |  |  |  |  |  |
| Asymptote ~ | Intercept | 147.81 | 2.97 | 49.80 | **<0.001\*\*\*** |
|  | Clutch size difference | 2.78 | 2.13 | 1.30 | 0.21 |
| Growth rate ~ | Intercept | 0.24 | 0.01 | 35.91 | **<0.001\*\*\*** |
|  | Clutch size difference | -0.01 | 0.01 | -1.37 | 0.19 |

## DISCUSSION

Cuckoo females should be under selection to maximise the quality of host parental care and choose individual hosts accordingly. By accounting for host availability and using clutch size as a proxy for individual quality, we predicted that common cuckoo females should select high-quality redstart nests when accounting for available host nests, and that cuckoo nestlings raised by chosen hosts should grow larger and/or faster. However, our results show that cuckoo females did not preferentially parasitise redstart nests with larger clutch sizes, despite clutch size being a proxy of individual quality in many passerines (Garamszegi et al., 2004; Houston et al., 1983). Nevertheless, experimentally disrupting individual host choice suggested that there could be weak but tantalising evidence that subverting the cuckoo femalés choice of individual host affected offspring growth. Our results therefore suggest that while clutch size may not be an ideal proxy for individual host quality, the choice of individual host nests could be adaptive for cuckoos.

### Is there scope for the cuckoo female to choose?

Our host nest availability analysis confirmed that there is variation (between 2 and 24) in the number of available nests between each theoretical cuckoo female breeding area of 800 m radius. Furthermore, during our spatio-temporal exploration of clutch size data, we found that redstart clutch sizes decreased throughout the season and increased in the north-western parts of our study area. This variation might be related to environmental differences (e.g., prey density, vegetation structure (Martinez, 2012; Tye, 1992)) or individual host differences, such as host female phenotype (Decker et al., 2012) or migration patterns (Bêty et al., 2003). Although we relied on estimating breeding areas of cuckoos around available nests, it is highly likely that these results captured host availability for four reasons. First, because natural crevices suitable for redstarts are scarce in our natural area (Moreras *et al*., 2021, pers. obs.). Second, because our nest boxes have very similar dimensions to those of natural cavities preferred by redstarts (Avilés et al., 2005), and thus redstarts readily accept our nest boxes (see Results). Third, because this situation allows us to accurately monitor redstart breeding cycle and record most redstart nests potentially available for cuckoo females. And four, because the few natural crevices that redstart might breed in are mostly woodpecker cavities, which are generally too small for the cuckoo female to lay her egg (Avilés et al., 2005; Moreras et al., 2021). Therefore, our results are strong evidence of quantitative and qualitative variation in potential host nest options for cuckoo females to parasitise. Notably, we obtained qualitatively similar results when repeating our analysis using three different putative cuckoo breeding area sizes (400 m, 800 m, and 1200 m radii).

Variation in host nest availability was further confirmed by variation in parasitism events in our area. We found that probability of parasitism showed spatio-temporal trends where parasitism increased throughout the breeding season and towards the south-west of the study area. Additionally, in our study area, cuckoo females parasitised hosts with different phenotypic traits, such as nests with clutches between 4 and 8 eggs. It has been shown that cuckoo females check the contents of several host nests before selecting one with their preferred phenotype, such as egg colour for instance (Zhang et al., 2023), or optimal laying timepoint during the host laying sequence (Wang et al., 2020). This suggests non-random choice (e.g., Soler *et al*., 1995; Wang *et al*., 2020) rather than a “shot-gun” laying approach (e.g., Orians, Røskaft and Beletsky, 1989; Lyon, 1993; Kattan, 1997). Therefore, our results confirmed that there is a range of choice for cuckoo females and that these parasites show variation in their choices. This could indicate that host nest selection could have an adaptive meaning in the Common cuckoo, even though we had to estimate the laying area of cuckoos rather than using genotyping to assign maternity.

### Do cuckoo females choose according to an individual quality proxy?

In our study, clutch size as a proxy for individual quality was not linked to a higher probability of parasitism, with several non-mutually exclusive explanations for this phenomenon. First, this could indicate that cuckoo females do not choose specific nests to lay their eggs. For instance, it has been shown that cuckoo females do not take into account individual host phenotype, such as egg shape and colour, when parasitising oriental reed warblers, *Acrocephalus orientalis* (Yang et al., 2016). Instead, cuckoo females could select nests at the appropriate breeding stage, i.e., still egg-laying (Wang et al., 2020). Second, it is possible that cuckoo females eavesdrop on visible and audible traits or behaviours, such as alarm calling during nest defence (Clotfelter, 1998; Marton et al., 2019) or within-pair communication (Parejo & Avilés, 2007). These cues are linked to parental quality in some species (Catchpole & Slater, 2003; Marton et al., 2019), but this remains unknown for common redstarts. Therefore, cuckoo females could be eavesdropping on redstartś alarm calls and songs because they might indicate individual quality or merely because they are easily detectable. Finally, it is possible that, even though clutch size has been traditionally described as a good quality proxy in other passerines (Garamszegi et al., 2004; Houston et al., 1983), it does not describe individual quality in common redstarts. Clutch size could instead be more influenced by e.g., vegetations structure (Martinez, 2012), changes in environmental conditions during the breeding season (Garamszegi et al., 2004), premigration conditions and migratory behaviour (Bêty et al., 2003), or even nest size (Soler et al., 2001) rather than by host female quality. To discriminate among these three potential explanations, further research is needed to determine which redstart traits are directly or indirectly linked to individual quality.

### What are the consequences of disrupting the cuckoo female choice?

Our cuckoo nestling growth results might be cautiously pointing towards a non-random egg-laying strategy. In our analysis of the effect of the type of nest on cuckoo nestling growth, we found similar weight and tarsus length growth between nestlings that were raised in their original nests and those that were moved to non-chosen nests. These results could indicate that cuckoo nestlings show a highly plastic behavioural adaptation to manipulate any individual host, mediated by a supernormal stimulus that compels foster parents to feed the parasitic nestling (Alvarez, 2004). In fact, it has been previously shown that cuckoo nestlings tune their begging depending on the species they parasitise (Madden &

Davies, 2006), and thus, this could also be happening at an individual host level. However, we found that cuckoo nestlings that were raised in their original nest reached a higher asymptotic growth of wings (mean ± SE = 8.22 ± 3.95 mm longer, Table 4) at a slower growth rate than nestlings moved to a non-chosen nest, with similar results in our exploratory analysis using clutch size difference as the predictor variable.

Growth performance with high asymptote and low growth rate or *vice versa* has been previously described in species with highly asynchronous hatching. In these nests, advantaged nestlings can take longer to reach a higher asymptotic growth, while disadvantaged nestlings need to grow faster at the cost of final size or mass (e.g., Zárybnická *et al*., 2015). Even though our results should be interpreted with caution due to the small sample size (n = 18), these could indicate that in advantageous growth environments (i.e. growing in nests selected by their mothers) cuckoo nestlings match their foster parents to be better able to extract care. This could be mediated by a quality match between cuckoo female and foster parent. Good quality hosts arrive early to the breeding grounds and thus select high quality areas (Decker et al., 2012). Similarly, high quality cuckoo females could also arrive earlier after the migration and select high quality hosts. If cuckoo nestlings inherit their motheŕs quality, then nestling quality should match host quality, therefore matching the cuckoo nestling begging behaviour to the host parents provisioning behaviour, as occurs in some other avian species (Estramil et al., 2013). This maternally mediated strategy would allow cuckoo nestlings to extract as much care as possible from their foster parents and optimise their growth. However, to clarify this potential mechanism, replicating our cross-fostering experiment with a larger sample size would be necessary. Additionally, we need further research on cuckoo female quality to increase our understanding of brood parasite- host interactions.

## CONCLUSION

The results of the present study indicate that there is scope for cuckoo female host choice. Our results also suggest that cuckoo females may select specific redstart hosts, and that those choices have consequences for nestling growth. However, both our results of cuckoo female choice and cuckoo nestling growth are determined by our estimate of the extension of cuckoo female breeding areas. This estimate was calculated based on recent work using molecular methods and egg phenotype to determine cuckoo maternal identity of cuckoo eggs and nestlings parasitising *Acrocephalus* warbler nests (Koleček et al., 2021). Nevertheless, cuckoo female breeding areas vary between studies using different hosts (Vogl *et al*., 2004; Nakamura, Miyazawa and Kashiwagi, 2005; Williams *et al*., 2016; Moskát *et al*., 2019). Therefore, to accurately understand cuckoo female host choice in our system and its consequences it is necessary to have actual measurements of cuckoo breeding areas when parasitising common redstarts. Alternatively, Koleček *et al*. could replicate our analysis testing whether clutch size as an individual quality proxy predicts parasitism probability when accounting for spatio- temporal host nest availability. Their experimental measurements of cuckoo female breeding areas when parasitising *Acrocephalus* warbler hosts could provide the perfect opportunity to accurately account for host nests available to each specific cuckoo female and further our understanding of host choice in brood parasitism.

## Supporting information

Supplementary material - Abaurrea et al 2024

## ACKNOWLEDGEMENTS

We thank all the field assistants for their help during fieldwork. We also thank Jesús Abaurrea and Jesús Asín (University of Zaragoza, Spain), as well as the Biodata Analytics Unit (University of Helsinki, Finland) for statistical advice. This study was supported by the Finnish Cultural Foundation and the Doctoral salaried positions of the University of Helsinki, both granted to T.A., and by several other funding institutions throughout the years. None of the funders had any influence on the content of the submitted or published manuscript.

## REFERENCES

1. Aggrey, S. (2002). Comparison of three nonlinear and spline regression models for describing chicken growth curves. Poultry Science, 81(12), 1782–1788. 10.1093/ps/81.12.1782

2. Alvarez, F. (2000). Clutches of rufous bush chats Cercotrichas galactotes parasitised by cuckoos Cuculus canorus contain larger eggs. Ardea, 88(1).

3. Alvarez, F. (2004). The conspicuous gape of the nestling Common cuckoo Cuculus canorus as a supernormal stimulus for Rufous bush chat Cercotrichas galactotes hosts. Ardea, 92(1), 63–68.

4. Avilés, J. M., Rutila, J., & Møller, A. P. (2005). Should the redstart Phoenicurus phoenicurus accept or reject cuckoo Cuculus canorus eggs? Behavioral Ecology and Sociobiology, 58(6), 608–617. 10.1007/s00265-005-0941-7

5. Banks, A. J., & Martin, T. E. (2001). Host activity and the risk of nest parasitism by brown-headed cowbirds. Behavioral Ecology, 12(1), 31–40. 10.1093/oxfordjournals.beheco.a000375

6. Bates, D., Mächler, M., Bolker, B., & Walker, S. (2014). *Fitting Linear Mixed-Effects Models using lme4* (arXiv:1406.5823). arXiv. 10.48550/arXiv.1406.5823

7. Bêty, J., Gauthier, G., & Giroux, J. (2003). Body Condition, Migration, and Timing of Reproduction in Snow Geese: A Test of the Condition-Dependent Model of Optimal Clutch Size. The American Naturalist, 162(1), 110–121. 10.1086/375680

8. Bryant, D. M. (1978). Environmental Influences on Growth and Survival of Nestling House Martins Delichon Urbica. Ibis, 120(3), 271–283. 10.1111/j.1474-919X.1978.tb06788.x

9. Catchpole, C. K., & Slater, P. J. B. (2003). Bird Song: Biological Themes and Variations. Cambridge University Press.

10. Clotfelter, E. D. (1998). What cues do brown-headed cowbirds use to locate red-winged blackbird host nests? Animal Behaviour, 55(5), 1181–1189. 10.1006/anbe.1997.0638

11. Davies, N. B. (2000). Cuckoos, Cowbirds and Other Cheats. T. & A.D. Poyser.

12. Decker, K. L., Conway, C. J., & Fontaine, J. J. (2012). Nest predation, food, and female age explain seasonal declines in clutch size. Evolutionary Ecology, 26(3), 683–699. 10.1007/s10682-011-9521-7

13. Estramil, N., Eens, M., & Müller, W. (2013). Coadaptation of Offspring Begging and Parental Provisioning—An Evolutionary Ecological Perspective on Avian Family Life. PLOS ONE, 8(7), e70463. 10.1371/journal.pone.0070463

14. Fossøy, F., Sorenson, M. D., Liang, W., Ekrem, T., Moksnes, A., Møller, A. P., Rutila, J., Røskaft, E., Takasu, F., Yang, C., & Stokke, B. G. (2016). Ancient origin and maternal inheritance of blue cuckoo eggs. Nature Communications, 7(1), Article 1. 10.1038/ncomms10272

15. Gao, C. Q., Yang, J. X., Chen, M. X., Yan, H. C., & Wang, X. Q. (2016). Growth curves and age-related changes in carcass characteristics, organs, serum parameters, and intestinal transporter gene expression in domestic pigeon (Columba livia). Poultry Science, 95(4), 867–877. 10.3382/ps/pev443

16. Garamszegi, L. Z., Török, J., Tóth, L., & Michl, G. (2004). Effect of timing and female quality on clutch size in the Collared Flycatcher Ficedula albicollis. Bird Study, 51(3), 270–277. 10.1080/00063650409461363

17. Geltsch, N., Hauber, M. E., Anderson, M. G., Bán, M., & Moskát, C. (2012). Competition with a host nestling for parental provisioning imposes recoverable costs on parasitic cuckoo chick’s growth. Behavioural Processes, 90(3), 378–383. 10.1016/j.beproc.2012.04.002

18. Gladbach, A., Gladbach, D. J., & Quillfeldt, P. (2010). Seasonal clutch size decline and individual variation in the timing of breeding are related to female body condition in a non-migratory species, the Upland Goose Chloephaga picta leucoptera. Journal of Ornithology, 151(4), 817– 825. 10.1007/S10336-010-0518-8/FIGURES/4

19. Grim, T. (2006). Cuckoo growth performance in parasitized and unused hosts: Not only host size matters. Behavioral Ecology and Sociobiology, 60(5), 716–723. 10.1007/s00265-006-0215-z

20. Grim, T., & Samaš, P. (2016). Growth Performance of Nestling Cuckoos Cuculus canorus in Cavity Nesting Hosts. Acta Ornithologica, 51(2), 175–188. 10.3161/00016454AO2016.51.2.004

21. Grim, T., Samaš, P., Procházka, P., & Rutila, J. (2014). Are tits really unsuitable hosts for the Common Cuckoo? Ornis Fennica, 91, 166–177.

22. Houston, D. C., Jones, P. J., & Sinly, R. M. (1983). The effect of female body condition on egg laying in Lesser black-backed gulls Larus fuscus. Journal of Zoology, 200(4), 509–520. 10.1111/j.1469-7998.1983.tb02812.x

23. Hussell, D. J. T. (1972). Factors Affecting Clutch Size in Arctic Passerines. Ecological Monographs, 42(3), 317–364. 10.2307/1942213

24. Jelínek, V., Procházka, P., Požgayová, M., & Honza, M. (2014). Common Cuckoos Cuculus canorus change their nest-searching strategy according to the number of available host nests. Ibis, 156(1), 189–197. 10.1111/ibi.12093

25. Kattan, G. H. (1997). Shiny cowbirds follow the ‘shotgun’ strategy of brood parasitism. Animal Behaviour, 53(3), 647–654. 10.1006/anbe.1996.0339

26. Koleček, J., Piálková, R., Piálek, L., Šulc, M., Hughes, A. E., Brlík, V., Procházka, P., Požgayová, M., Capek, M., Sosnovcová, K., Štětková, G., Valterová, R., & Honza, M. (2021). Spatiotemporal patterns of egg laying in the common cuckoo. Animal Behaviour, 177, 107–116. 10.1016/j.anbehav.2021.04.021

27. Louder, M. I. M., Schelsky, W. M., Albores, A. N., & Hoover, J. P. (2015). A generalist brood parasite modifies use of a host in response to reproductive success. Proceedings of the Royal Society B: Biological Sciences, 282(1814), 20151615. 10.1098/rspb.2015.1615

28. Lyon, B. E. (1993). Tactics of parasitic American coots: Host choice and the pattern of egg dispersion among host nests. Behavioral Ecology and Sociobiology, 33(2), 87–100. 10.1007/BF00171660

29. Madden, J. R., & Davies, N. B. (2006). A host-race difference in begging calls of nestling cuckoos Cuculus canorus develops through experience and increases host provisioning. Proceedings of the Royal Society B: Biological Sciences, 273(1599), 2343–2351. 10.1098/rspb.2006.3585

30. Marshall, D. J., & Uller, T. (2007). When is a maternal effect adaptive? Oikos, 116(12), 1957–1963. 10.1111/j.2007.0030-1299.16203.x

31. Martinez, N. (2012). Sparse vegetation predicts clutch size in Common Redstarts Phoenicurus phoenicurus. Bird Study, 59(3), 315–319. 10.1080/00063657.2012.672949

32. Marton, A., Fülöp, A., Ozogány, K., Moskát, C., & Bán, M. (2019). Host alarm calls attract the unwanted attention of the brood parasitic common cuckoo. Scientific Reports, 9(1), Article 1. 10.1038/s41598-019-54909-1

33. Molina-Morales, M., Martínez, J. G., & Avilés, J. M. (2016). Criteria for host selection in a brood parasite vary depending on parasitism rate. Behavioral Ecology, 27(5), 1441–1448. 10.1093/beheco/arw066

34. Moreras, A., Tolvanen, J., Morosinotto, C., Bussiere, E., Forsman, J., & Thomson, R. L. (2021). Choice of nest attributes as a frontline defense against brood parasitism. Behavioral Ecology, 32(6), 1285–1295. 10.1093/beheco/arab095

35. Moskát, C., Bán, M., Fülöp, A., Bereczki, J., & Hauber, M. E. (2019). Bimodal habitat use in brood parasitic Common Cuckoos (Cuculus canorus) revealed by GPS telemetry. The Auk, 136(2), uky019. 10.1093/auk/uky019

36. Nakamura, H., & Miyazawa, Y. (1997). Movements, Space Use and Social Organization of Radio- tracked Common Cuckoos during the Breeeding Season in Japan. Japanese Journal of Ornithology, 46, 23–54.

37. Nakamura, H., Miyazawa, Y., & Kashiwagi, K. (2005). Behavior of radio-tracked Common Cuckoo females during the breeding season in Japan. Ornithological Science, 4(1), 31–41. 10.2326/osj.4.31

38. Orians, G. H., Røskaft, E., & Beletsky, L. D. (1989). Do Brown-Headed Cowbirds Lay Their Eggs at Random in the Nests of Red-Winged Blackbirds? The Wilson Bulletin, 101(4), 599–605.

39. Pagani-Núñez, E., & Senar, J. C. (2014). Are colorful males of great tits Parus major better parents? Parental investment is a matter of quality. Acta Oecologica, 55, 23–28. 10.1016/j.actao.2013.11.001

40. Parejo, D., & Avilés, J. M. (2007). Do avian brood parasites eavesdrop on heterospecific sexual signals revealing host quality? A review of the evidence. Animal Cognition, 10(2), 81–88. 10.1007/s10071-006-0055-2

41. Polačiková, L., Procházka, P., Cherry, M. I., & Honza, M. (2009). Choosing suitable hosts: Common cuckoos Cuculus canorus parasitize great reed warblers Acrocephalus arundinaceus of high quality. Evolutionary Ecology, 23(6), 879–891. 10.1007/s10682-008-9278-9

42. R Core Team. (2023). *R: A language and environment for statistical computing.* [Computer software]. http://www.R-project.org/

43. Ricklefs, R. E. (1968). Patterns of Growth in Birds. Ibis, 110(4), 419–451. 10.1111/j.1474-919X.1968.tb00058.x

44. Rutila, J., Latja, R., & Koskela, K. (2002). The common cuckoo Cuculus canorus and its cavity nesting host, the redstart Phoenicurus phoenicurus: A peculiar cuckoo-host system? Journal of Avian Biology, 33(4), 414–419. 10.1034/j.1600-048X.2002.02937.x

45. Schwagmeyer, P. L., & Mock, D. W. (2008). Parental provisioning and offspring fitness: Size matters. Animal Behaviour, 75(1), 291–298. 10.1016/j.anbehav.2007.05.023

46. Silva, M. C., Boersma, P. D., Mackay, S., & Strange, I. (2007). Egg size and parental quality in thin- billed prions, Pachyptila belcheri: Effects on offspring fitness. Animal Behaviour, 74(5), 1403– 1412. 10.1016/j.anbehav.2007.01.008

47. Slagsvold, T., & Lifjeld, J. T. (1990). Influence of Male and Female Quality on Clutch Size in Tits (Parus Spp.). Ecology, 71(4), 1258–1266. 10.2307/1938263

48. Soler, J. J., Neve, L. de, Martínez, J. G., & Soler, M. (2001). Nest size affects clutch size and the start of incubation in magpies: An experimental study. Behavioral Ecology, 12(3), 301–307. 10.1093/beheco/12.3.301

49. Soler, J. J., Soler, M., Møller, A. P., & Martínez, J. G. (1995). Does the great spotted cuckoo choose magpie hosts according to their parenting ability? Behavioral Ecology and Sociobiology, 36(3), 201–206. 10.1007/BF00177797

50. Starck, J. M., & Ricklefs, R. E. (1998). Avian Growth and Development: Evolution Within the Altricial- precocial Spectrum. Oxford University Press.

51. Stokke, B. G., Hafstad, I., Rudolfsen, G., Moksnes, A., Møller, A. P., Røskaft, E., & Soler, M. (2008). Predictors of resistance to brood parasitism within and among reed warbler populations. Behavioral Ecology, 19(3), 612–620. 10.1093/beheco/arn007

52. Thomson, R. L., Tolvanen, J., & Forsman, J. T. (2016). Cuckoo parasitism in a cavity nesting host: Near absent egg-rejection in a northern redstart population under heavy apparent (but low effective) brood parasitism. Journal of Avian Biology, 47(3), 363–370. 10.1111/jav.00915

53. Trnka, A., Požgayová, M., Procházka, P., Prokop, P., & Honza, M. (2012). Breeding success of a brood parasite is associated with social mating status of its host. Behavioral Ecology and Sociobiology, 66(8), 1187–1194. 10.1007/s00265-012-1372-x

54. Tye, A. (1992). Assessment of territory quality and its effects on breeding success in a migrant passerine, the Wheatear Oenanthe oenanthe. Ibis, 134(3), 273–285. 10.1111/j.1474-919X.1992.tb03810.x

55. Vogl, W., Taborsky, B., Taborsky, M., Teuschl, Y., & Honza, M. (2004). Habitat and space use of European cuckoo females during the egg laying period. Behaviour, 141(7), 881–898. 10.1163/1568539042265671

56. Wang, L., Yang, C., He, G., Liang, W., & Møller, A. P. (2020). Cuckoos use host egg number to choose host nests for parasitism. Proceedings of the Royal Society B: Biological Sciences, 287(1928), 20200343. 10.1098/rspb.2020.0343

57. Wilkin, T. A., King, L. E., & Sheldon, B. C. (2009). Habitat quality, nestling diet, and provisioning behaviour in great tits Parus major. Journal of Avian Biology, 40(2), 135–145. 10.1111/j.1600-048X.2009.04362.x

58. Williams, H. M., Willemoes, M., Klaassen, R. H. G., Strandberg, R., & Thorup, K. (2016). Common Cuckoo home ranges are larger in the breeding season than in the non-breeding season and in regions of sparse forest cover. Journal of Ornithology, 157(2), 461–469. 10.1007/s10336-015-1308-0

59. Winnicki, S. K., Strausberger, B. M., Antonson, N. D., Burhans, D. E., Lock, J., Kilpatrick, A. M., & Hauber, M. E. (2021). Developmental asynchrony and host species identity predict variability in nestling growth of an obligate brood parasite: A test of the “growth-tuning” hypothesis. Canadian Journal of Zoology, 99(3), 213–220. 10.1139/cjz-2020-0147

60. Wood, S. N. (2017). Generalized Additive Models: An Introduction with R, Second Edition. CRC Press.

61. Yang, C., Wang, L., Liang, W., & Møller, A. P. (2016). Egg recognition as antiparasitism defence in hosts does not select for laying of matching eggs in parasitic cuckoos. Animal Behaviour, 122, 177–181. 10.1016/j.anbehav.2016.10.018

62. Zárybnická, M., Riegert, J., Brejšková, L., Šindelář, J., Kouba, M., Hanel, J., Popelková, A., Menclová, P., Tomášek, V., & Šťastný, K. (2015). Factors Affecting Growth of Tengmalm’s Owl (Aegolius funereus) Nestlings: Prey Abundance, Sex and Hatching Order. PLOS ONE, 10(10), e0138177. 10.1371/journal.pone.0138177

63. Zhang, J., Santema, P., Lin, Z., Yang, L., Liu, M., Li, J., Deng, W., & Kempenaers, B. (2023). Differences in the costs and benefits of choosiness may explain variation in cuckoo egg-matching strategy: A reply to Wang and Liang (2023). Proceedings of the Royal Society B: Biological Sciences, 290(2006), 20231219. 10.1098/rspb.2023.1219

64. Zuur, A. F. (2014). *A beginner’s guide to generalised additive mixed models with R.* Highland Statistics Ltd.

