## Supplementary material - Abaurrea et al 2024 for "Common cuckoo female host selection is not determined by host quality but can affect cuckoo nestling growth when parasitising Common redstarts"

Results of spatio-temporal variation in redstart clutch size using a putative cuckoo female breeding area of smaller (400 m) and larger (1200 m) radius than the one estimated during our host nest availability analysis (800 m).

**Results of analysis using a 400 m radius cuckoo breeding area**.

Table SM1: Model selection of parasitism GLMMs.

| *Model* | *K* | *AIC* | *Delta AIC* | *AIC weight* |
|---|---|---|---|---|
| ~ STA clutch size + Available nests + (1\|Year) | 4 | 716.56 | 0.00 | 0.73 |
| ~ STA clutch size * Available nests + (1\|Year) | 5 | 718.52 | 1.95 | 0.27 |
| ~ 1 + (1\|Year) | 2 | 729.61 | 13.04 | 0.00 |
| ~ STA clutch size + (1\|Year) | 3 | 730.71 | 14.15 | 0.00 |

Table SM2. GLMM explaining the parasitism probability of redstart nests according to spatio-temporally adjusted (STA) clutch size. Number of available nests is included as a covariate to control for variation in potential choices. SE: standard error. Significant p-values in bold (*** , P<0.001; ** , P<0.01; * , P<0.05; . , P<0.1).

| Response: Probability of parasitism | Estimate | SE | z-value | p-value |
|---|---|---|---|---|
| Intercept | -0.43 | 0.28 | -1.55 | 0.12 |
| STA clutch size | -0.09 | 0.11 | -0.85 | 0.40 |
| Available nests | -0.27 | 0.07 | -3.82 | **<0.001*** ** |

**Results of analysis using a 1200 m radius cuckoo breeding area**.

Table SM3: Model selection of parasitism GLMMs.

| *Model* | *K* | *AIC* | *Delta AIC* | *AIC weight* |
|---|---|---|---|---|
| ~ STA clutch size + Available nests + (1\|Year) | 4 | 872.57 | 0.00 | 0.71 |
| ~ STA clutch size * Available nests + (1\|Year) | 5 | 874.32 | 1.75 | 0.29 |
| ~ 1 + (1\|Year) | 2 | 908.89 | 36.31 | 0.00 |
| ~ STA clutch size + (1\|Year) | 3 | 910.14 | 37.57 | 0.00 |

Table SM4. GLMM explaining the parasitism probability of redstart nests according to spatio-temporally adjusted (STA) clutch size. Number of available nests is included as a covariate to control for variation in potential choices. SE: standard error. Significant p-values in bold (*** , P<0.001; ** , P<0.01; * , P<0.05; . , P<0.1).

| Response: Probability of parasitism | Estimate | SE | z-value | p-value |
| --- | --- | --- | --- | --- |
| Intercept | -0.48 | 0.20 | -2.38 | **0.017 *** |
| STA clutch size | -0.09 | 0.10 | -0.91 | 0.36 |
| Available nests | -0.07 | 0.01 | -5.88 | **<0.001 ***** |
